# A novel deep learning pipeline for cell typing and phenotypic marker quantification in multiplex imaging

**DOI:** 10.1101/2022.11.09.515776

**Authors:** Ettai Markovits, Tal Dankovich, Roman Gluskin, Ido Weiss, Amit Gutwillig, Tomer Dicker, Sun Dagan, Ron Elran, Becky Arbiv, Yuval Shachaf, Amit Bart, Assaf Debby, Nethanel Asher, Guy Ben-Betzalel, Ronnie Shapira-Frommer, Iris Barshack, Ori Zelichov

**Affiliations:** Nucleai, Tel Aviv, Israel; Department of Pathology, Sheba Medical Center, Ramat Gan, Israel; Ella Lemelbaum Institute of Immuno-Oncology, Sheba Medical Center, Ramat Gan, Israel; Tel-Aviv University, Tel Aviv, Israel

## Abstract

**Background:** Multiplex immunofluorescence (mIF) can provide invaluable insights into spatial biology and the complexities of the immune tumor microenvironment (iTME). However, existing analysis approaches are both laborious and highly user-dependent. In order to overcome these limitations we developed a novel, end-to-end deep learning (DL) pipeline for rapid and accurate analysis of both tumor-microarray (TMA) and whole slide mIF images.

**Methods:** Our pipeline consists of two DL models: a multi-classifier for classifying multi-channel cell images into 12 different cell types, and a binary classifier for determining the positivity of a given marker in single-channel images. The DL multi-classifier was trained on 7,000 tiles labeled with cell annotations from a publicly available CODEX dataset, consisting of 140 tissue cores from 35 colorectal cancer (CRC) patients. For the binary classifier training, the multi-channel tiles were further split into ∼100,000 single-channel tiles, for which the ground truth was inferred from the known expression of these markers in each cell-type. This DL binary classifier was then utilized to quantify the positivity of various cell state (phenotypic) markers. In addition, the binary classifier was exploited as a cell-typing tool, by predicting the positivity of individual lineage cell markers. The performance of our DL models was evaluated on 1,800 annotations from 14 test tissue cores. The models were further evaluated on a new 6-plex melanoma cohort, stained with PhenoImager®, and were compared to the performance of clustering, manual thresholding or machine learning-based cell-typing methods applied on the same test sets.

**Results:** Our DL multi-classifier achieved highly accurate results, outperforming all of the tested cell-typing methods, including clustering, manual-thresholding and ML-based approaches, in both CODEX CRC and PhenoImager melanoma cohorts (accuracy of 91% and 87%, respectively), with F1-scores above 80% in the vast majority of cell types. Our DL binary classifier, which was trained solely on the lineage markers of the CRC dataset, also outperformed existing methods, demonstrating excellent F1-scores (>80%) for determining the positivity of unseen phenotypic and lineage markers across the two tumor types and imaging modalities. Notably, as little as 20 annotations were required in order to boost the performance on an unseen dataset to above 85% accuracy and 80% F1-scores. As a result, the DL binary classifier could successfully be used as a cell-typing model, in a manner that is transferable between experimental approaches.

**Conclusions:** We present a novel state-of-the-art DL-based framework for multiplex imaging analysis, that enables accurate cell typing and phenotypic marker quantification, which is robust across markers, tumor indications, and imaging modalities.

## Introduction

Cellular organization within tissues is a crucial aspect for understanding the biological functions and processes in health, disease and malignancy^1^. Spatial biology provides invaluable insights into the complex interactions and relationships between the distinct cells that drive tissue functions in health, and underlie the orchestration of immune response against pathogens and tumors^2^. Accordingly, recognition has grown for the importance of the tumor microenvironment (TME) and immune composition within the tumor area in determining the clinical outcomes of immunotherapy-treated cancer patients^3,4^. As a result, several tools have been developed to extract high-dimensional cellular properties while preserving tissue-wide spatial context, at the forefront of which are multiplexed tissue imaging technologies such as multiplex immunofluorescence (mIF).

In a mIF experiment, iterative staining, imaging, and washing cycles are applied to acquire up to 100 unique markers, which are then mapped to distinct cells within a single histological section, providing a highly-detailed roadmap of the TME^5,6^. However, the analysis of mIF is hindered by the challenge in combining data from marker intensities to meaningful biological features, such as cell types and cell states of identified cells. As current existing methods such as clustering and manual thresholding, are laborious, user-dependent and underperforming, especially in rare cell types^7,8^, there is a need for a more robust and automated pipeline for multiplex imaging analysis. Recently, an unsupervised machine learning algorithm for automated cell typing of multiplex images was developed^9^, which reported F1-scores between 0.6 and 0.7 across major cell types and between 0.4 and 0.6 for rare cell types using a CODEX colorectal cancer (CRC) dataset. However, the ground truth used for calculating these metrics was clustering-based cell typing which validity is not yet established.

Here, we present a novel end-to-end DL pipeline to analyze mIF images, allowing for rapid and accurate analysis of multiplex imaging across markers, tumor indications and imaging modalities.

## Results

### A deep learning-based framework for cell typing in multiplex imaging

Multiplex imaging is a powerful tool for understanding the role of spatial biology, tissue architecture and cell-cell interactions in the pathophysiology of different diseases and response to treatments^6, 10^. Typically, multiplex imaging analysis begins with the identification of cells, which are further classified into different cell types using clustering- or manual thresholding-based methods, based on the cells’ expression of various lineage markers. Next, phenotypic markers are quantified as either continuous or binary variables for each cell, usually by averaging marker expression and setting a manual threshold for each marker. These methods are laborious, user-dependent, and do not support transfer learning between imaging modalities and antibody panels^5, 11^. Moreover, the accuracy and performance of these models in cell typing and marker quantification are seldom evaluated and reported. To overcome these limitations, we developed a DL-based framework for cell typing and marker quantification based on annotations by expert annotators (Figure 1A). To assess the framework performance relative to current analysis methods, we leveraged a publicly available multiplex immunofluorescence dataset^7^, consisting of 140 tissue cores from 35 colorectal cancer (CRC) patients stained with 56 protein markers, to annotate 12 cell types, as well as positivity for 14 immunomodulatory proteins (Figure 1B). We utilized over 7,000 cell annotations (100-1,000 annotations per class; Figure S1A) on 58 tissue cores to train a DL multi-channel classifier (DL multi-classifier) which classifies cells into 12 subpopulations. For training, cells’ nuclei were segmented and 20×20um^2^ multi-channel tiles containing both the lineage markers and the segmentation mask were cropped around each annotated cell. The tiles were divided into training and validation datasets, and a test set of 1,800 annotations from 14 test cores, which the model was not trained upon. Our DL multi-classifier reached a 92% accuracy (88.8%-99.6% balanced accuracy per class), with over 80% of cell classes demonstrating above 85% F1-scores in the test tissue cores (Figure 1C, D). To compare our results to existing clustering-based methods, we mapped between cell classes from *Schurch et al*.^7^ and our own annotations (Table S1A). The clustering-based cell-typing approach achieved only 63.4% balanced accuracy, with less than 10% of cell classes reaching F1-scores above 85% (Figure 1D). Next, we compared our cell typing results to a ML-based multiclass classifier which was trained on the same data. Although the overall accuracy of the ML-based algorithm was 88% (Figure S1B) the F1-scores of B-cells (96% vs. 64%) and dendritic cells (74% vs. 35%) were much lower in the ML-based multiclass classifier (Figure S1C). Taken together, we demonstrate that our novel DL pipeline for multiplex imaging cell typing, exhibits high accuracy, and a major improvement over current cell-typing methods.

**Figure 1.**
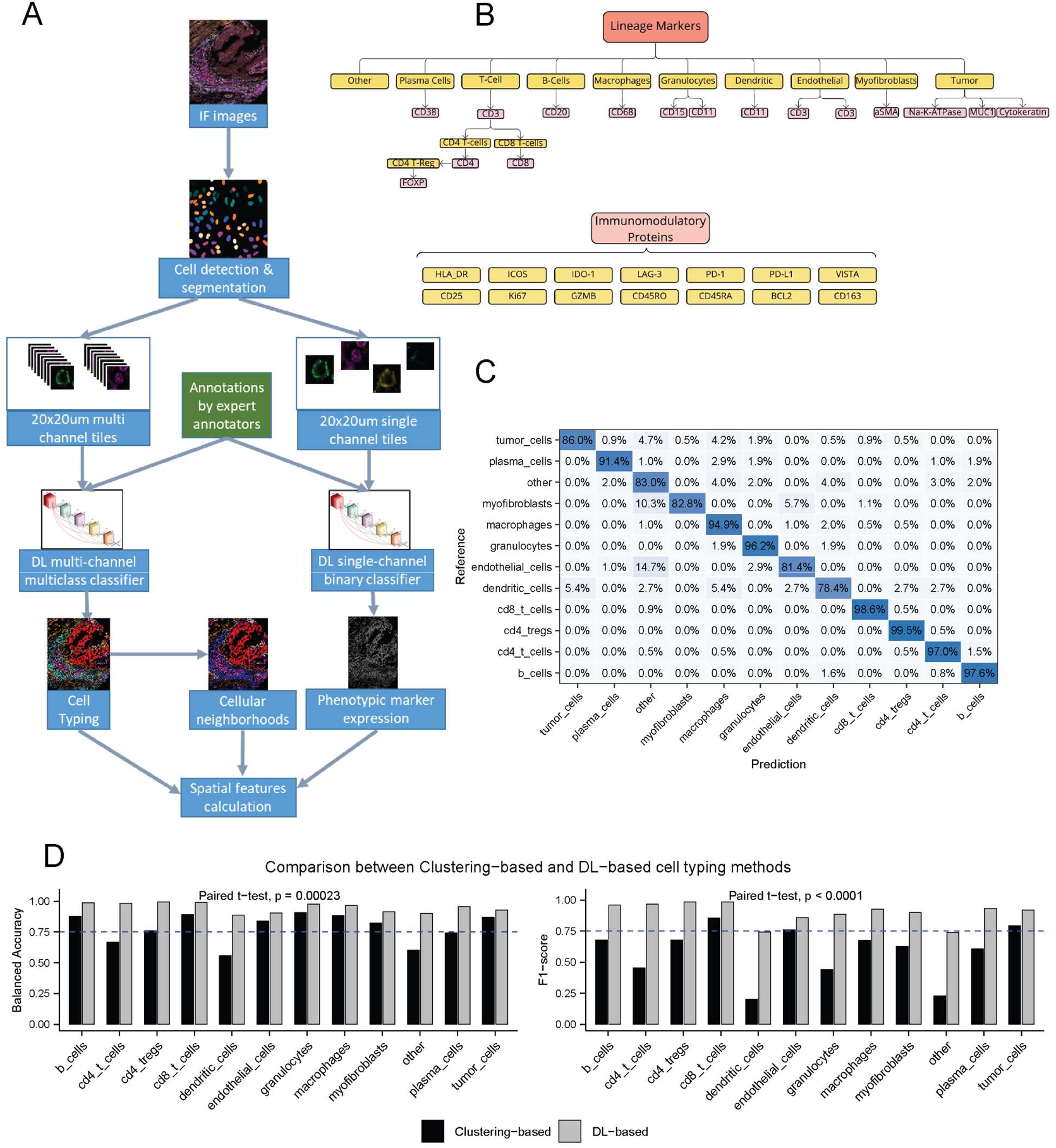
A novel deep-learning (DL) pipeline for cell-typing in multiplex imaging. **A**, Schematic of the analysis pipeline. mIF images containing a panel of cell lineage markers are fed into a DL-based cell segmenter to identify all cell instances. A training set of segmented cells is annotated with cell types by expert annotators, which are then cropped into 20×20 µm^2^ tiles and fed into a DL-based multi- or binary classifier. For the former (left side of the panel), the tiles are fed into the model as multi-channel images, and then classified into one of the annotated cell types. For the latter model (right side of the panel), the tiles are first split into single-channel images, whose ground truth is based on the expected lineage marker expression based on reports in the literature. The output (positivity/negativity of an individual fluorescence channel), is utilized to predict the positivity of previously unseen fluorescence channels of phenotypic markers, and determine their expression in the identified cells. The cell types and phenotypic marker expression can then be utilized to calculate a diversity of spatial features that may be used to predict clinical outcomes. **B**, Depiction of the lineage and immunomodulatory (phenotypic) markers stained in the publicly available dataset that was utilized in this study (C. M. Schürch et al., 2020). **C**, Confusion matrix for the DL multi-classifier predicted cell types. The model achieved an overall accuracy of 92% (88.8%-99.6% balanced accuracy per class). **D**, A comparison of the balanced accuracy (left) and F1 scores (right) between our DL multi-classifier and a clustering-based cell-typing approach. The DL-based approach outperformed clustering for both metrics across all cell types (paired *t*-test; *p* = 0.0023 and *p* < 0.0001 for the comparison of balanced accuracy (left) and F1 scores (right) between the clustering- and DL-based approaches).

### A deep learning-based single-channel binary quantifier for phenotypic markers quantification

Besides the utilization of high-plex data for identification of cellular subpopulations, a major advantage of multiplex imaging over current immunohistochemistry and IF imaging methods is that it allows to deduce cell functionality from phenotypic marker expression. Existing methods for the deduction of phenotypic marker positivity include either continuous quantification from segmented cell masks, which is influenced by both the segmentation quality and the stain intensity variation between slides, or manual thresholding, which is user-dependent, laborious, and is not robust between slides. Thus, there is a need for a model that can classify cell markers in a binary fashion, that will be robust across slides, markers, tissue type and imaging modalities. To develop our DL-based single-channel binary classifier, we utilized known marker expression (Table S1B) to divide the 7,000 annotated multi-channel tiles, used to train the DL cell classifier, into more than 90,000 binary single-channel annotated tiles (Figure 2A). The DL binary classifier outputs a positive prediction probability for each tile, which is then thresholded to output a binary classification. As a baseline threshold, we used 0.5 probability to classify tiles as either negative or positive. The model was evaluated on 19,000 single-channel lineage marker tiles in the 14 test tissue cores and exhibited excellent performance, reaching >90% overall accuracy and F1-score (Figure 2B). However, per class examination of the F1-scores revealed F1-scores below 75% in 18% of lineage markers (3/17). Indeed, further inspection of the positive prediction probability distribution in positive and negative tiles demonstrated uneven distributions between markers and slides (Figure 2C, S2). Thus, we performed per-channel threshold optimization on the training set (PCTO), choosing thresholds which maximize F1-scores, while taking into account positive prediction probability variability between slides. While we observed only a mild increase in overall F1 scores following per-channel threshold optimization to 92%, the overall accuracy increased to 97% and both CD31 and Na-K-ATPase demonstrated a significant performance boost in F1 scores after PTCO (76% vs. 89% and 77% vs. 87%, respectively, Figure 2B). Next, we evaluated the binary classifier model performance on 14 phenotypic markers in 14 test tissue cores, which the binary classifier was not trained upon. We evaluated the model performance on 1,600 single-channel phenotypic marker annotations (40-200 annotations per marker) and, using 0.5 as a threshold, observed a 91% accuracy, with F1 scores of above 80% in 93% of markers (Figure 2D). Per channel threshold optimization with annotations from the train set tissue cores boosted the overall F1-score to 95%, which stemmed from a significant improvement in the F1-scores of markers with relatively low positive prediction probabilities such as LAG3 (89% to 98%), HLA-DR (82% to 97%) and PD-L1 (62% to 92%, Figure 2D-E). To investigate the required number of annotations per channel for threshold optimization, we compared the F1-scores of the 5 markers which improved with PTCO with increasing numbers of annotations. Remarkably, only 20 annotations per channel were needed to achieve the most significant improvement in F1-scores, with additional annotations adding limited value (Figure 2F). As the DL binary classifier is robust across markers and slides, it could be used to accurately quantify a wide variety of cell state and lineage markers with a minimal number of added annotations to improve its performance in specific channels.

**Figure 2.**
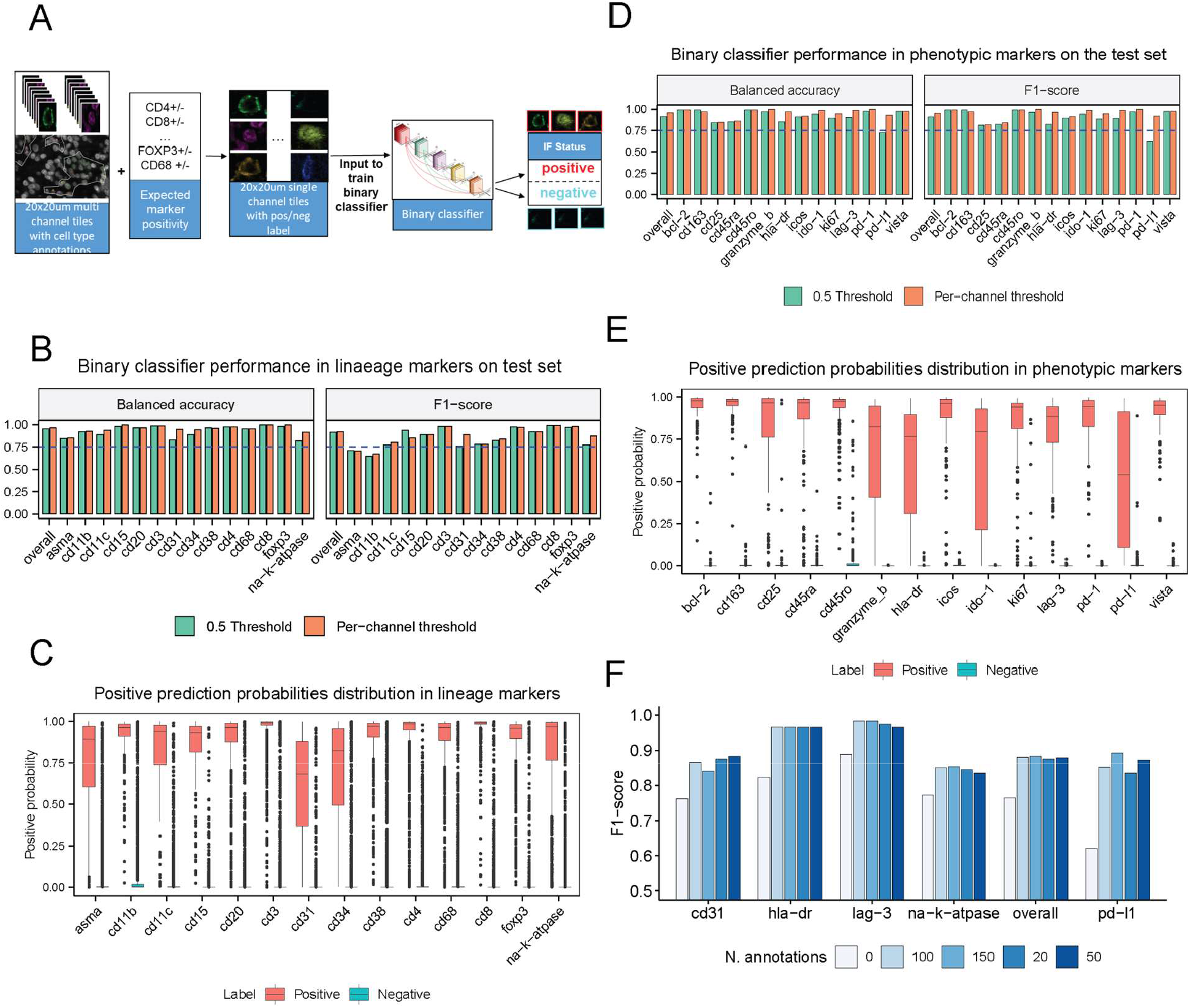
A single-channel binary classifier can be utilized to predict the positivity of cell state markers. **A**, The 7000 multi-channel tiles annotated with 12 different cell types (used to train the DL multi-classifier) were further split into >90,000 single-channel tiles representing lineage markers. Literature-based knowledge of lineage marker expression within these cell types (Table S1B) was used to establish the ground truth (positive/negative expression) for the tiles. **B**, The model was evaluated on 19,000 single-channel lineage marker tiles in 14 test tissue cores. With a global threshold of 0.5 for predicting positive expression, the classifier reaches >90% overall accuracy and F1-score. Improved results are achieved with per channel threshold optimization (PCTO). **C**, The distribution of the probability of positivity across lineage markers, for tiles with positive (red) and negative (blue) single-channel tiles. Although the distribution is expected to differ significantly between positive and negative tiles, the result is highly variable across cell types. **D**, The DL binary classifier was evaluated on 1,600 annotated tiles of 14 phenotypic markers (40-200 annotations per marker) which the model had not been trained on. Using a default threshold of 0.5, the model achieved 91% accuracy, with F1 scores >80% in 93% of markers. The F1 score was improved to 95% when applying PCTO. **E**, The distribution of the probability of positivity across phenotypic markers, for tiles with positive (red) and negative (blue) single-channel tiles. **F**, The F1 scores achieved for DL binary classifier prediction of various lineage cell markers using PCTO with 20, 50, 100 or 150 annotations, as compared to a default threshold of 0.5 (labeled ‘0 annotations’ in the plot). Only 20 annotations are required in order to obtain a significant improvement in the F1 score.

### Utilization of the DL single-channel binary classifier for ‘label free’ cell typing

The high performance of the DL single-channel classifier across lineage markers raises the question whether it can be utilized as a ‘label-free’ cell-typing tool. To create ‘auto-labels’, we combined our knowledge of expected marker expression within cell classes (Table S1B) with the DL binary classifier single-channel predictions and positive probabilities for each marker (Figure 3A). Cells were assigned to a class if their binary marker predictions perfectly matched the expected binary marker expression of this class. Next, we created a table of the mean positive probability for each marker within each cell class and then matched between unassigned cells and their closest mean positive probability vector. Comparison between auto-labels generated with the 0.5 threshold for marker positivity (‘label free’) and annotated cells within the test tissue cores, demonstrated an overall good performance with a 88.1% accuracy and F1-scores above 80% in 75% of cell classes. Using PTCO with 20 annotations for low performing channels (CD31, CD11c, αSMA and Na-K-ATPase), we were able to increase the model’s performance to 91% accuracy (Figure 3B) while increasing the F1-scores for tumor (84.8% vs 90.5%), myofibroblasts (81.5% vs. 88%) and endothelial cells (86% vs. 91.5%) (Figure 3C). Thus, our single-channel model reached a level of performance that was comparable to the multi-classifier (Figure 3C), with the exception of two cell classes; dendritic cells and other, in which the F1-scores were lower.

**Figure 3.**
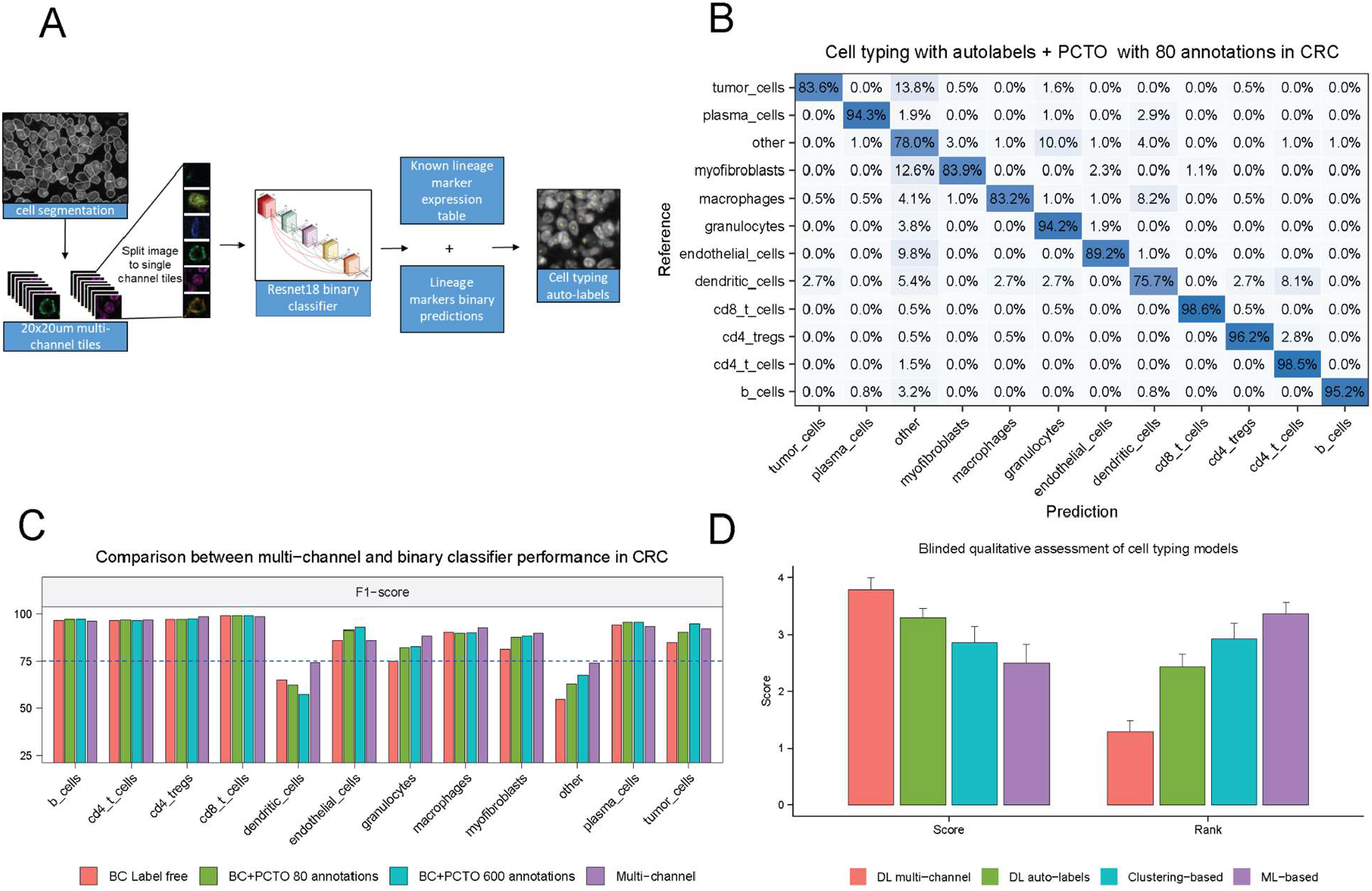
Utilization of the DL binary classifier as a cell-typing model. **A**, To convert the single-channel output of the DL binary classifier into cell types (‘auto-labels’), we combined the lineage marker predictions from the binary classifier with expected marker expression per cell type, as reported in the literature. **B**, Confusion matrix of the ‘auto-labels’ predicted by the DL binary classifier on the CRC test set, using per-channel threshold optimization (PCTO) with 20 annotations per channel to optimize the binary prediction threshold. An overall accuracy of 91% was achieved. **C**, A comparison of the F1 scores between the DL multi-classifier and the DL binary classifier with PCTO using 80 or 600 annotations in total (for all channels). With the exception of the ‘dendritic cells’ and ‘other’ classes, the binary classifier achieved results that are comparable to the multi-classifier, with just 80 annotations. **D**, A qualitative comparison between the clustering-, ML-based, DL multi-classifier and DL binary classifier-based ‘auto-labels’ models, based on ranking by expert annotators. The DL multi-classifier was ranked highest, with the DL binary-classifier ‘auto-labels’ ranking second.

To further corroborate our model’s performance, we qualitatively compared between the four cell typing models: clustering-based, ML-based, DL multi-classifier, and ‘auto-labels’ from the DL binary classifier. The models were blindly ranked and scored by expert annotators, based on their performance on the 14 tissue cores. The DL multi-classifier was the top performing model, as reflected by 86% first-place votes and overall performance score of 3.8/5 (Figure 3D). The second-best model was ‘auto-labels’ from the DL binary classifier, which received the majority of second-place votes, and an overall performance score of 3.3. Both the ML and clustering-based algorithms received a performance score that was lower than 3, and were only ranked as the top two performing models in 25% of the cases (Figure 3D). Thus, we demonstrate that DL-based cell-typing is superior to current approaches, and present a pipeline for cell-typing through single-channel predictions which reaches comparable results to that of the DL multi-classifier, but requires significantly less annotations.

### Validation of the DL binary classifier performance across markers, cancer indication and imaging modalities

One of the main limitations of multiplex imaging analysis is that all current analysis methods do not allow for transfer learning between datasets. We demonstrated that the DL binary classifier is robust across markers, including markers it was not trained upon, and can be used for cell typing. However, it is yet to be proven whether our models will perform well on a different imaging platform and tumor indication. To establish the robustness of the DL binary classifier across different antibodies, tumor types and imaging modalities we stained 43 melanoma whole slide FFPE sections for 6 markers (CD8, CD4, FOXP3, CD68, SOX10 and PD-L1) with PhenoImager technology (Figure 4A). The images were divided to train (n=35) and test (n=9) sets. The binary classifier, which was trained on the CRC codex dataset, was deployed for ‘label free’ cell-typing to predict marker positivity for the lineage markers and deduce auto-labels (Figure 4A). Without any annotations, the binary classifier performed well on all lineage markers, reaching F1-scores above 80% in all markers (Figure S3A). However, a closer look at the positive prediction probabilities revealed lower positive prediction scores for SOX10 (Figure S3B). Following threshold optimization for SOX10 using only 20 annotations, the F1-score of SOX10 increased from 81% to 90%. In comparison to manual-thresholding, the current state-of-the-art method for establishing marker positivity, the binary classifier exhibited a better overall F1-score (85.7% vs. 78.5%), as well as better F1-scores in 80% of quantified markers (Figure 4B). Hence, the binary classifier is robust across tumor indication and imaging modalities, outperforming current state-of-the-are methods for marker positivity determination.

**Figure 4.**
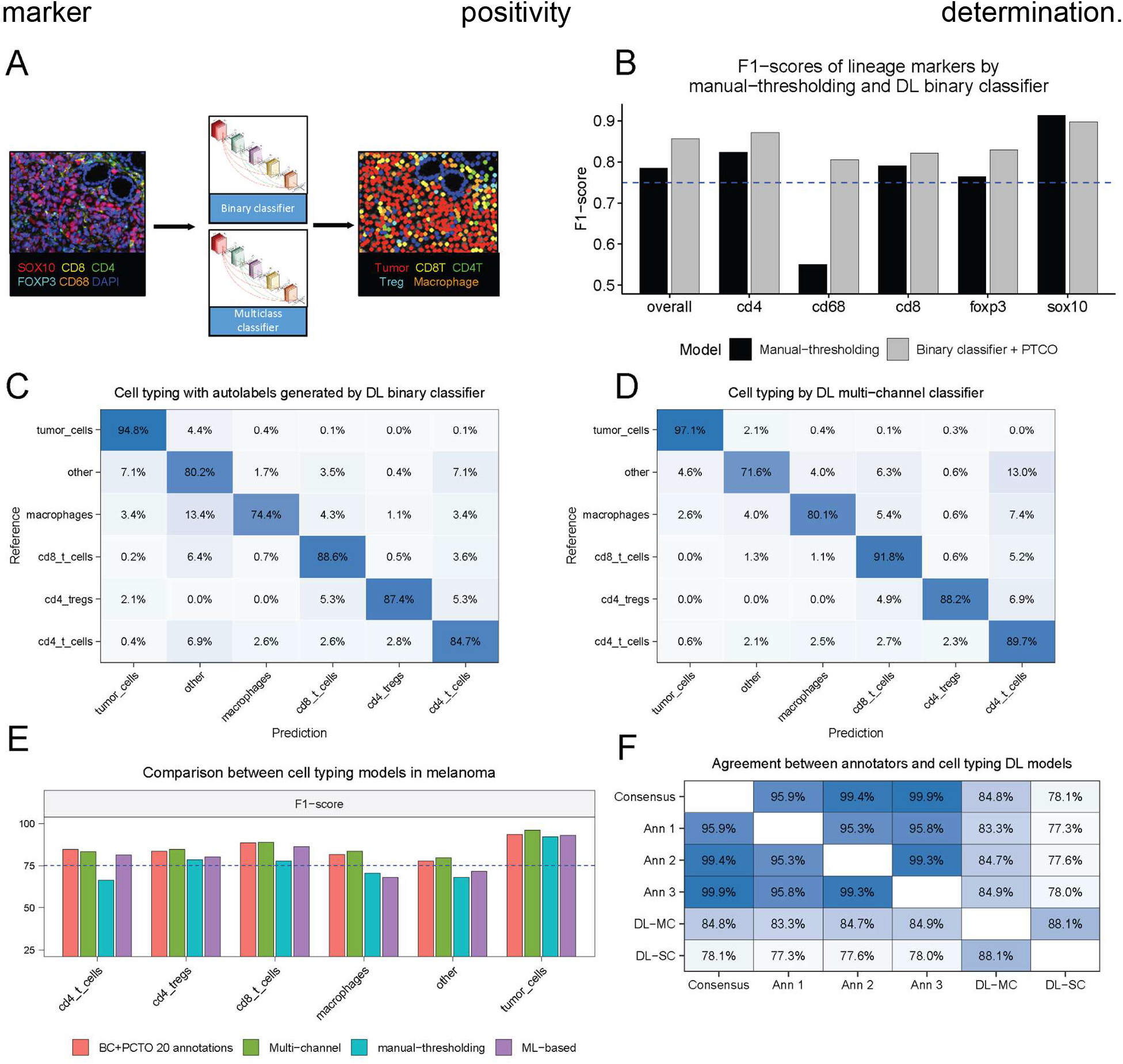
Application of the DL binary classifier for cell-typing on a new tumor type and imaging modality. **A**, 43 melanoma WSIs were stained with a panel of 6 markers with PhenoImager® technology. The DL binary classifier, trained on lineage markers of the CRC dataset, was deployed to create ‘auto-labels’ for 6 different classes in the unseen melanoma WSIs, based on our knowledge of linear marker expression. The DL multi-classifier was trained to classify cells on 21,000 annotations from the melanoma dataset. **B**, A comparison of the F1 scores for the prediction of previously unseen lineage markers by the DL binary classifier with per-channel threshold optimization (PCTO), or by a manual thresholding. The binary classifier exhibited a better overall F1-score (85.7% vs. 78.5%), as well as better F1-scores in 80% of quantified markers. **C**, Confusion matrix of the ‘auto-labels’ predicted by the DL binary classifier for 6 different classes on unseen data. An 86% overall accuracy was achieved using this approach. **D**, Confusion matrix of a DL multi-classifier trained on the melanoma PhenoImager data. The overall accuracy was 87%. **E**, A comparison of F1 scores between the DL multi-classifier (trained on the current dataset), DL binary classifier (trained on the CRC dataset), manual thresholding-based approach, and an ML-based classifier (trained on the current dataset) across classes. Lower F1 scores are observed for the ML and manual thresholding-based algorithms. **F**, The inter-observer agreement rate between expert annotators, and comparison to model predictions. Three experts independently annotated the same 1,200 cells within the melanoma PhenoImager dataset to establish the validity of our annotations. This was further compared to the predictions of the DL multi-classifier and the DL binary classifier ‘auto-labels’ (trained on the CRC dataset). >95% agreement was observed between the annotators, and an 85% and 75% agreement with the annotation consensus was observed for the DL multi-classifier and binary classifiers, respectively.

Next, we compared our ‘label free’ cell-typing based on ‘auto-labels’ generated by the binary classifier or manual thresholding, with current state-of-the-art ML and DL methods for multiclass classification. Both a DL multi-classifier and a ML-multi-classifier were trained on over 15,000 cell type annotations from 35 training set slides (Figure S3C). All models were evaluated on 2,800 annotations from 9 test set slides. Cell typing by our DL-based models were superior to both the ML- and manual thresholding-based models, as reflected by higher accuracy (87%, and 86% for the DL multi- and binary classifier (Figure 4C, D) vs. 81% and 79% for the ML and manual-thresholding based classifiers, (Figure S3D, E). Moreover, a comparison of per-class F1-scores demonstrated similar F1-scores between the DL-based algorithm, with lower F1-scores in the ML and manual thresholding based algorithms (Figure 4E). Lastly, we aimed to compare our models’ performance with the inter-observer agreement rate between expert annotators. For this purpose, three annotators labeled a total of 1200 cells within identical exhaustive ROIs in 5 WSIs from the melanoma cohort. We observed >95% inter-observer agreement rate between all three annotators, establishing the validity of annotations as ground truth for multiplex imaging. The DL multi-classifier exhibited an 85% agreement with the annotation consensus (defined as agreement between two annotators or more) while the ‘auto-labels’ generated by the DL binary classifier exhibited a 78% agreement with the consensus (Figure 4F). Thus, our multiplex analysis pipeline demonstrated robustness across markers, tumor type and imaging modalities and can be deployed for a rapid and accurate analysis of multiplex imaging. While the DL multi-classifier vastly surpassed current methodologies performance for cell typing, establishing new state-of-the-art performance benchmark, the DL binary classifier can be utilized for phenotypic marker quantification and demonstrated only a slight reduction in performance in comparison to the DL multi-classifier, while relying on a miniscule number of annotations.

## Discussion

The spatial organization of cells in the iTME has an essential role in the process of tumor formation and immune system evasion. mIF imaging of tissue biopsies using recently developed tools, such as CODEX, has emerged as a powerful tool for iTME analysis that could potentially be used to predict clinical outcomes in cancer patients, including response to immunotherapy and overall survival. However, the analysis of multiplex images suffers from several limitations including inaccurate cell-typing, which is mainly achieved through manual thresholding and clustering-based methods. These methods are laborious, user-dependent, and do not support transfer learning between imaging modalities and antibody panels, which makes them hard to implement as a routine tool for translational and clinical research. To overcome these challenges we developed a deep learning pipeline for the analysis of mIF images that can be generalized across tissue types, markers and imaging modalities.

Our pipeline utilizes both multi-channel and single-channel deep learning classifiers and achieves a high accuracy of over 90% in classifying cells, as compared to ∼65-80% using manual thresholding, clustering or machine learning-based cell-typing methods. One of the features that makes this pipeline unique is our binary single-channel classifier which is agnostic to marker type and can accurately classify markers that it was not trained upon. Thus, it can be utilized for both phenotypic marker classification and cell typing, as it identifies cell classes by relying on expected marker expression while using minimal annotations. As a result, this novel mIF analysis pipeline is significantly faster and more robust across markers and slides, as compared with other methods. Indeed, the model reached an overall accuracy of >95% both in classifying lineage markers that it was trained on and in classifying phenotypic markers which it was not trained upon. More importantly, the DL binary classifier demonstrated robustness across different antibodies, tumor types and imaging modalities - while the binary classifier was trained on CODEX CRC dataset, it showed high accuracy (86%) when tested on melanoma sections stained with PhenoImager imaging technology, with minimal addition of annotations. Lastly, we demonstrate a very high inter-observer agreement rate between expert annotators (>95%), which validates that annotations should be used as ground truth for multiplex imaging. The DL-multi-classifier exhibited high agreement rate with the annotators consensus (85%), again establishing it as state-of-the-art benchmark for cell typing. Taken together, we exhibit a novel DL pipeline for multiplex imaging cell typing, demonstrating high accuracy and a 1.5-fold improvement over current cell typing methods, and robustness across markers, tumor type and imaging modalities. Thus, it can potentially be used for a rapid and accurate analysis of multiplex imaging.

## Materials and Methods

### Datasets and annotations

For model generation, two multiplex imaging datasets were used. The first is a publicly available CODEX dataset^7^, consisting of 140 tissue cores from 35 colorectal cancer (CRC) patients, stained with 56 protein markers and matched H&E slides. The images were downloaded and annotated by expert annotators, under the supervision of expert pathologists. Above 7,000 cell annotations from 57 tissue cores were used as a training set for training of the DL and ML classifiers. 1,800 annotations from 14 test cores, which the models were not trained upon, were used for performance evaluation of the models. Moreover, over 1600 annotations of positive and negative cells for 14 phenotypic markers (Figure 1A) on 14 test cores were used as a test set for the DL binary classifier performance. A second cohort, consisting of 44 whole slide images (WSIs) of melanoma cancer patients from Sheba medical center, was stained with 6 protein markers (Figure 4A) and captured with PhenoImager® imaging system (Akoya Biosciences). Over 15,000 cell type annotations from 35 training set slides were used as a training set, while 2,800 annotations from 9 test set slides were used for model evaluation.

### Cell segmentation

Multi-instance cell segmentation was performed using a deep learning model. Nuclear segmentation was based on the DAPI and Hoechst channels (for the CRC and melanoma datasets, respectively), and whole cell segmentation was based on the sum of all available membranous and cytoplasmic channels. Further post-processing was performed to match nuclear and whole cell masks, allowing to remove segmented cells that do not contain nuclei, merge the nuclei of multinucleated cells and split any nuclei assigned to multiple cells. In addition, a membranal segmentation mask was extracted by subtracting each nuclear mask from its matched whole cell mask, and then regularizing it by a ring around the nucleus.

### Convolutional deep neural networks for cell typing and phenotypic markers quantification

As input for the model 54×54-pixel (20 µm^2^) tiles were cropped around the centerpoint of all annotated cell segmentation instances. The tiles were then normalized to and scaled between 0 and 1. All models were trained based on Nucleai Ltd deep learning infrastructure, and were trained for at least 500 epochs. For the DL-multi-classifier, the 54×54-pixel tiles were fed into the model as multi-channel images. For the CRC dataset, multi-channel tiles consisting of 15 lineage markers (Figure 1B) were inputted to the model, whereas for the melanoma dataset multi-channel tiles consisting of 5 lineage markers (Figure 4A) were inputted. For the single-channel classifier (DL binary classifier), the tiles were split into single-channel images, and the ground truth per channel was determined via the known marker expression table (Table S1B). Both the multiclass and binary classifiers output prediction probabilities. For the multi-classifier, prediction probabilities are converted to cell types by assigning the cell type with maximal prediction probability to each identified cell. For the binary classifier, cells above a certain threshold are deemed positive. The threshold can be set to 0.5 for all channels or adjusted based on positive or negative tile annotations (per-channel thresholding optimization; PTCO). For subsequent cell-typing from single-channel binary classification of lineage markers (‘auto-label’ deduction), the expected marker expression tables (Table S1B, C) were applied to the thresholded channels.

### Clustering-based cell typing

In order to compare between our cell typing results to the clustering based algorithm deployed in *Schurch et al*.^7^, we mapped between our 12 identified cell types and the 28 identified clusters described by *Schurch et al*. (Table S1A). Clusters were mapped by the maximal representative class present in our training set annotations, where clusters with less than 10 matching cells were ignored, and classes with the maximal fraction of a representative class of less than 0.25 were assigned to ‘other’.

### XGboost classifier for cell typing

The segmentation masks of cells were used to calculate the mean nuclear and membranous fluorescence intensity per channel for all annotated cells instances in the CRC dataset. These features were then Yeo-Johnson normalized to approximate normal distribution and scaled between 0 and 1. A multiclass cell XGBoost classifier was hyperparameter optimized using a cross-validated randomized search. Each sample was assigned a weight based on the square root of the class incidence (with the rare dendritic cell upweighted by 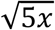).

### Cell typing by manual thresholding of melanoma images with HALO®

For cell-typing by manual thresholding, 5 ROIs in 34 WSI from the melanoma dataset were spectrally unmixed using inForm® software v6.4.2 (Akoya Biosciences), and analyzed with the HALO® image analysis platform (Indica Labs), using the Highplex FL module. In addition, spectral auto-fluorescence reduction was performed from an unstained but rehydrated slide which was scanned under identical conditions. Following identification and segmentation of ∼665,000 cells, the positivity for each marker was determined by an algorithm that was trained on the positive control tonsil sample to identify the intensity threshold within the appropriate cell compartment. All tissue samples were examined to ensure that the algorithm identified each marker accurately, and in some cases, a manual adjustment was required to optimize the thresholding of specific channels.

### Model performance evaluation

All models were evaluated on test slides that were not seen during training. We considered annotations by expert annotators as the ground truth, and performance metrics such as accuracy, balanced accuracy and F1-scores were calculated for each model.To qualitatively evaluate the models trained on the CRC CODEX dataset, a thorough review was performed under the guidance of experienced pathologists, and each model was given a score between 1-5, per WSI. The scoring was based on the percentage of overall cell detection for each model: 0-20, 20-40, 40-60, 60-80, 80-100 (1-5 levels). If a cell class was over- or under-detected, a point was deducted from the overall score. Additionally, all models were ranked from top to bottom according to their performance in each slide.

### Agreement test

To assess the consistency of our cell annotations and model performance, we conducted an independent agreement test between three expert annotators. The test set consisted of ∼1200 cells per annotator, from exhaustive ROIs in 5 melanoma WSIs images. The consensus cell-typing was decided by the majority vote of annotators.

## Supplementary Figures

**Figure S1, related to Figure 1.**
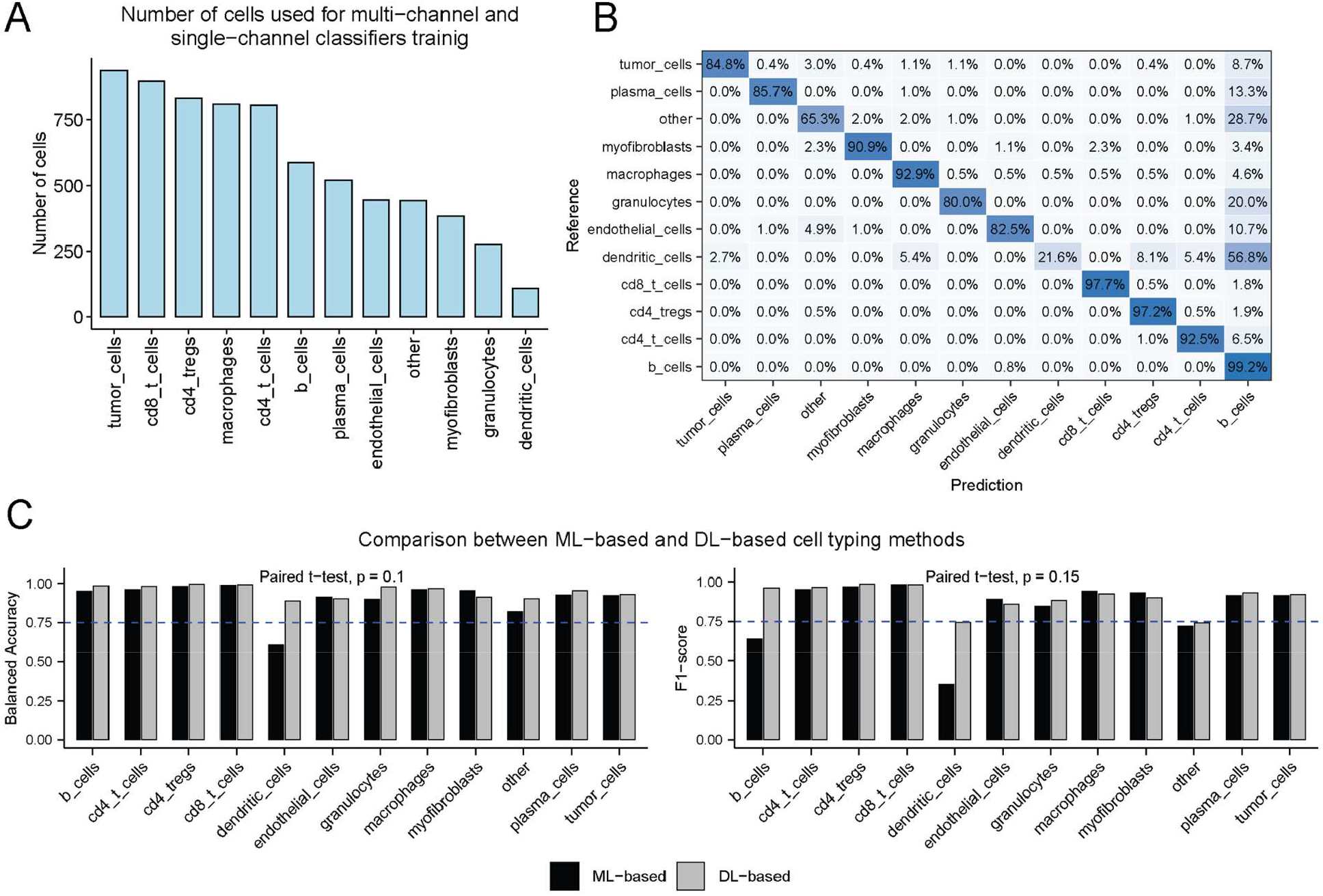
**A**, Breakdown of the number of cell type annotations on the CRC CODEX dataset for each of the 12 classes. **B**, Confusion matrix for the ML-based multi-classifier (XGBoost), which reached an overall accuracy of 88%. **C**, A comparison of the balanced accuracy (top) and F1 scores (bottom) between our DL multi-classifier and an ML-based cell-typing approach. While the overall performance of of the DL-based classifier was not significantly different, the F1-scores of B-cells (96% vs. 64%) and dendritic cells (74% vs. 35%) were higher for the DL-based approach (paired *t*-test; *p* = 0.1 and *p* = 0.15 for the comparison of balanced accuracy and F1 scores, respectively).

**Figure S2. Related to figure 2.**
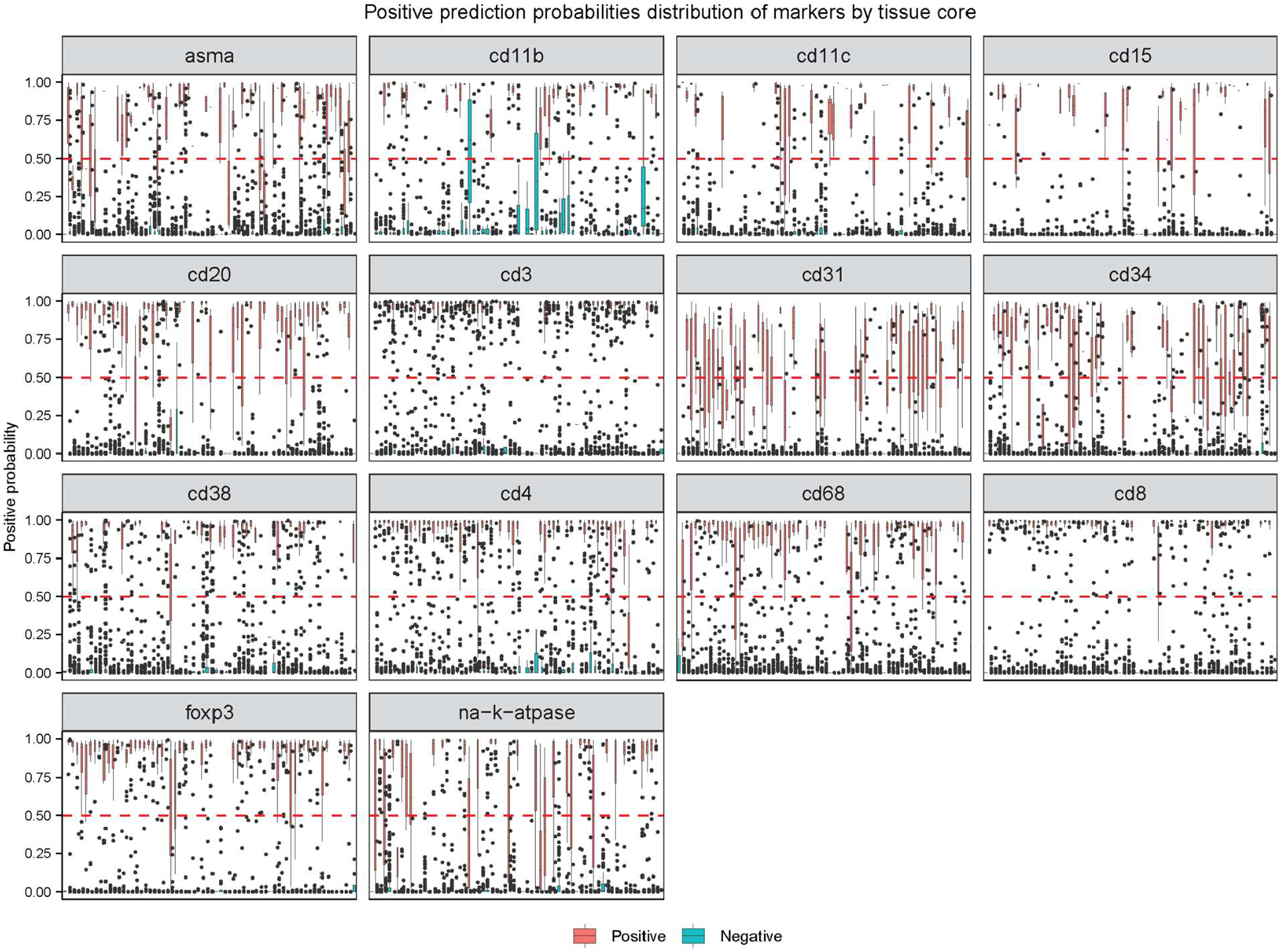
The distribution of the probability of positivity across lineage markers, for tiles with positive (red) and negative (blue) single-channel tiles, for different tissue cores.

**Figure S3. Related to Figure 4.**
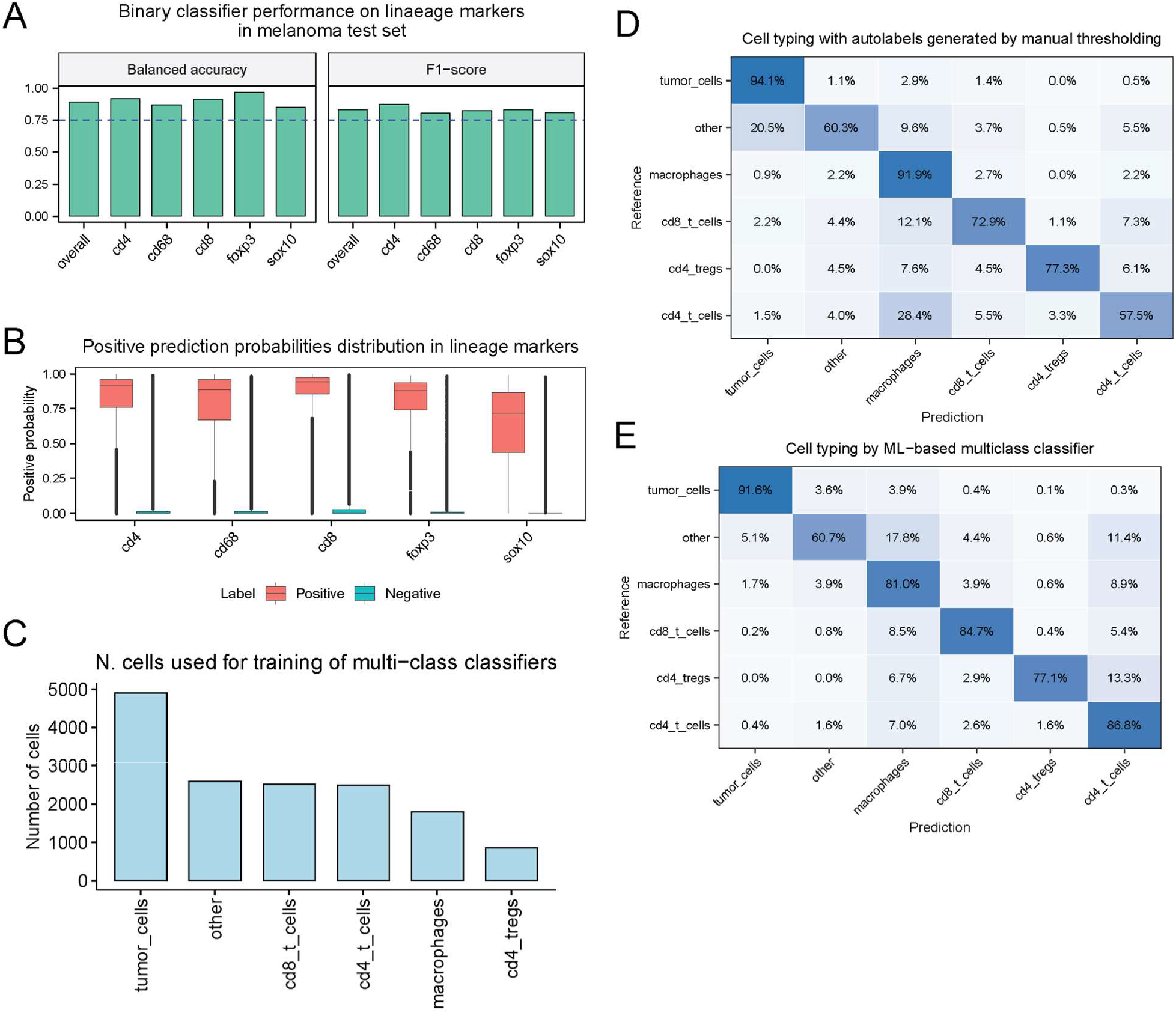
**A**, Performance of the DL binary classifier (balanced accuracy and F1 scores), trained on the CRC dataset on lineage markers from an unseen melanoma dataset, across all lineage markers. In the absence of any annotations, the binary classifier performed well, reaching F1-scores >80% for all markers. **B**, The distribution of the probability of positivity across lineage markers, for tiles with positive (red) and negative (blue) single-channel tiles. It can be seen that the difference between the positive/negative distributions is less apparent for SOX10, where lower positive prediction scores are observed. **C**, Breakdown of the number of cell type annotations on the melanoma PhenoImager dataset for each of the 6 classes. **D**, Confusion matrix of the cell-typing produced by a manual thresholding approach, considered to be the current state-of-the-art (see Methods), which achieved an overall accuracy of 81%. **E**, Confusion matrix fan ML-based multi-classifier (XGBoost) trained on the melanoma PhenoImager dataset, which reached an overall accuracy of 79%.

